# Exploring the PLD1-tau interaction in Frontotemporal Dementia

**DOI:** 10.64898/2026.02.12.705569

**Authors:** Chandramouli Natarajan, Shaneilahi M. Budhwani, Sravan Gopalkrishna Shetty Sreenivasamurthy, Laharika Katamoni, Bernadette Thomson, Michela Marcatti, Phan Phu Cuong, Giulio Taglialatela, Balaji Krishnan

## Abstract

**Summary:** Frontotemporal dementia (FTD), a leading cause of young-onset dementia, is characterized by progressive behavioral and cognitive decline associated with frontotemporal cortical atrophy. Nearly 40% of cases exhibit tauopathy, yet the molecular drivers of tau aggregation leading to synaptic dysfunction remain poorly understood. Here, we investigated whether Phospholipase D1 (PLD1, a lipid signaling enzyme), implicated in Alzheimer’s disease (AD), and amyotrophic lateral sclerosis (ALS), contributes to tau pathology dependent synaptic deficits in FTD.

Postmortem temporal (BA38) and frontal (BA9) cortices from clinically diagnosed FTD and age-matched control subjects were analyzed using fluorescence-assisted single synaptosome long-term potentiation (FASS-LTP), immunofluorescence, proximity ligation assays (PLA), and PLD1-interactome proteomics.

FASS-LTP revealed markedly reduced glutamatergic potentiation in BA38 and BA9 crude synaptoneurosomes from FTD brains compared to controls. Western blotting demonstrated elevated PLD1 expression in both crude synaptoneurosomal and cytosolic fractions from FTD subjects in BA38, but not BA9. Bielschowsky staining confirmed increased Pick body burden in FTD temporal cortex. Immunofluorescence and PLA showed robust PLD1 co-localization with total tau (HT7), hyperphosphorylated tau (AT8), and acetylated tau oligomers (TOMA2), indicating a strong spatial association between PLD1 and pathological tau species. PLD1 also exhibited enhanced co-localization with astrocytic GFAP and synaptic markers (PSD95, Nrx1β), suggesting compartmentalized involvement in glial and synaptic remodeling. Proteomic profiling of PLD1-associated complexes revealed compartment-specific alterations with cytosolic fractions enriched for metabolic enzymes, stress-response proteins, and GFAP, while crude synaptoneurosomal fractions showed depletion of presynaptic scaffolds, vesicle-trafficking regulators, and proteostasis components. Cross-compartment integration indicated that over one-third of proteins were redistributed from synapses to cytosol, consistent with trafficking and degradative impairments. Gene Ontology analysis highlighted lipid metabolism, astrocyte activation, and proteasome dysfunction as dominant pathways.

Collectively, these findings identify PLD1 as a critical mediator of synaptic dysfunction and tau pathology in FTD, acting through astroglial activation and disrupting synaptic proteostasis. This study provides the human clinical relevance towards PLD1 attenuation as a therapeutic target for FTD and related tauopathies to mitigate tau-driven neurodegeneration and restore synaptic integrity.

## Introduction

Frontotemporal dementia (FTD) is the second most common cause of young-onset dementia, presenting with progressive behavioral, language, and cognitive impairments driven by frontal and temporal lobe degeneration[1–7]. Approximately 40% of FTD cases are classified as primary tauopathies, characterized by pathological tau aggregation, neuronal loss, gliosis, and the presence of argyrophilic inclusions known as Pick bodies[8–11]. While memory deficits are less pronounced early in FTD compared to Alzheimer’s disease (AD), synaptic dysfunction is increasingly recognized as a central mechanism underlying cognitive decline across tauopathies[12,13].

Phospholipase D1 (PLD1), a member of the PLD superfamily of lipolytic enzymes, hydrolyzes phosphatidylcholine to generate phosphatidic acid—a lipid mediator critical for membrane curvature, vesicle trafficking, and exocytosis[14–18]. PLD1 signaling influences synaptic vesicle release and receptor trafficking, processes essential for maintaining synaptic integrity[17,18]. Previous work from our group has implicated PLD1 in tau oligomer-driven synaptic impairment and memory deficits in AD models and human AD brains, establishing PLD1 as a key node in lipid-mediated synaptic dysfunction[19–23]. Furthermore, PLD1 has emerged as a therapeutic target in related neurodegenerative conditions such as amyotrophic lateral sclerosis (ALS), where its downregulation showed ameliorative effects[24,25]. ALS shares clinical and pathological overlap with FTD, reinforcing the need to investigate PLD1 in the ALS–FTD spectrum[13,26].

Despite these observations, the specific role of PLD1 and its interaction with pathological tau in FTD-related synaptopathy remains largely unexplored. We hypothesized that PLD1 contributes to synaptic deficits observed in FTD through interactions with pathological tau and associated proteostatic disruptions. Quantitative neuropathological studies demonstrate that both postmortem anterior temporal (BA38) and dorsolateral prefrontal (BA9) cortices are among the first and most severely affected regions in FTD[27–30]. In a cohort of 38 FTD-MAPT autopsy cases, tau burden and neuronal degeneration were most severe for all tau isoforms in BA38, while tau pathological strains were significant in BA9, particularly in mixed isoform cases with consistently higher severity of neuronal degeneration[31,32]. Plasma lipidomic studies in FTD reveal region-specific alterations in multiple lipid classes, with particular relevance to the frontal and temporal cortices, in triglycerides with decreases in phospholipids (substrates for phospholipases including PLD1)[33]. Using a systematic strategy, we evaluated synaptic integrity in BA38 and BA9 through our validated fluorescence-assisted single-synaptosome long-term potentiation (FASS-LTP) assay as a functional measure[22,34–36]. Next, we quantified PLD1 expression in crude synaptoneurosomal and cytosolic fractions and evaluated its spatial association with tau species—including total tau, hyperphosphorylated tau (AT8), and acetylated tau oligomers (TOMA2)—using immunofluorescence (IF) and proximity ligation assays (PLA). We further investigated PLD1 co-localization with astrocytic (GFAP) and synaptic markers (PSD95, Nrx1β) to delineate compartment-specific involvement. Finally, PLD1-interactome proteomics was performed to identify protein networks associated with PLD1 in cytosolic and crude synaptoneurosomal fractions, enabling pathway-level insights into synaptic trafficking, proteostasis, and glial remodeling.

This integrated analysis demonstratesPLD1 dysregulation in synaptic dysfunction and tau pathology using human FTD brains, and revealing astrocytic contributions with proteomic signatures that underscore PLD1 as a potential therapeutic target for FTD and related tauopathies.

## Materials and Methods

### Human Postmortem Tissues

Postmortem temporal (BA38) and frontal (BA9) cortices were obtained from the NIH NeuroBioBank (Table 1).

**Table 1:** Demographic and Clinical Characteristics of Post-Mortem Human Brain Tissue Samples. This table presents list of de-identified post-mortem brain tissue samples obtained from the NeuroBioBank (NBB). The table includes requested information on clinical diagnosis (Frontotemporal Dementia - FTD, Lewy body dementia - LBD, or Control), post-mortem interval (PMI in hours), age (in years), and sex. <u>Key Observations:</u> *Sample Distribution:* The cohort includes eight control subjects, seven FTD subjects, and one LBD subject. *Age Range:* The age of the subjects ranges from 54 to 80 years. *Sex Distribution:* The cohort comprises 11 males and 5 females. *PMI:* from 7 to 37 hours.

| Subject ID | Clinical Diagnosis | PMI | Age | Sex |
| --- | --- | --- | --- | --- |
| **5080** | Control | 14 | 61 | Female |
| **5511** | Control | 10 | 80 | Female |
| **5597** | Control | 17 | 63 | Male |
| **5985** | Control | 23 | 55 | Male |
| **6065** | Control | 22 | 55 | Male |
| **5862** | Control | 26 | 58 | Male |
| **5890** | Control | 37 | 60 | Male |
| **5666** | Control | 25 | 65 | Male |
| **5632** | FTD | 20 | 63 | Female |
| **6149** | LBD | 7 | 76 | Female |
| **5151** | FTD | 12 | 65 | Male |
| **5729** | FTD | 13 | 60 | Male |
| **HBFW2** | FTD | 22.25 | 57 | Male |
| **6107** | FTD | 24 | 54 | Male |
| **6180** | FTD | 12 | 56 | Male |
| **HBER** | FTD | 20 | 64 | Male |

### Immunofluorescence

Fresh-frozen tissue blocks were equilibrated at –20 °C, embedded in O.C.T., and sectioned at 12–16 µm onto Superfrost Plus slides. Sections were fixed in 4% paraformaldehyde (PFA, pH 7.4) for 30 min at room temperature, blocked with 5% BSA and 10% normal goat serum, and permeabilized with 0.5% Triton X-100/0.05% Tween-20. Primary antibodies included PLD1, AT8, HT7, TOMA2, Neurexin-1β, PSD95, and GFAP (see Supplementary Table S1 for details). Slides were incubated overnight at 4 °C, washed, and treated with Alexa Fluor-conjugated secondary antibodies (1 h, RT). Autofluorescence was quenched with 0.3% Sudan Black in 70% ethanol, and slides were mounted with DAPI-containing Fluoromount-G. Representative images were acquired using a Keyence BZ-X810 microscope. Subjects with high PMI (5890) or limited tissue availability (5862 for PLD1–PSD95 analysis) were excluded. Detailed protocols, antibody catalog numbers, and RRIDs are provided in Supplemental Methods.

### Fluorescence-Assisted Single Synaptosome Long-Term Potentiation (FASS-LTP)

Crude synaptoneurosomal fractions (P2) from frontal (BA9) and temporal (BA38) cortices were prepared using the Syn-PER method[19]. FASS-LTP was performed as described previously[22,34–36]. Briefly, crude synaptoneurosomes were incubated in extracellular, basal, or chemical LTP (cLTP) solutions, followed by pharmacological stimulation with glycine and KCl in the presence of bicuculline and strychnine. After depolarization, crude synaptoneurosomes were labeled with antibodies against GluA1 and Neurexin-1β and analyzed by flow cytometry (Guava easyCyte™ 8). Double-positive events (GluA1⁺Nrx1β⁺) were quantified as a measure of synaptic potentiation. Detailed buffer compositions, incubation steps, and antibody specifications are provided in Supplemental Methods.

### Bielschowsky Staining

Silver staining was performed to visualize tau deposits and Pick bodies using a modified Bielschowsky protocol[37–39]. Briefly, cryosections were hydrated, impregnated with silver nitrate, developed, and toned with ammonium hydroxide. Stained sections were dehydrated and mounted for bright-field imaging. Detailed reagent compositions, incubation times, procedural steps, and quantitative analysis are provided in Supplemental Methods.

### Proximity Ligation Assay (PLA)

Protein proximity between PLD1 and hyperphosphorylated tau (AT8) was assessed using the Duolink® in situ PLA protocol (Sigma-Aldrich)[40]. Briefly, fixed temporal cortex sections from control and FTD subjects were incubated with primary antibodies against PLD1 and AT8, followed by PLUS/MINUS PLA probes, ligation, and rolling-circle amplification according to manufacturer’s instructions. Signals were visualized using fluorescence microscopy and quantified as described in Supplemental methods.

### Tissue Processing and Western Blot Analyses

Frontal (BA9) and temporal (BA38) cortices from FTD and control subjects were homogenized in Syn-PER buffer with protease/phosphatase inhibitors. Crude synaptoneurosomal (P2) and cytosolic (S2) fractions were prepared by differential centrifugation and resuspended in Krebs-like buffer. Protein concentrations were determined spectrophotometrically, resolved by SDS-PAGE, and transferred to nitrocellulose membranes. Blots were probed with antibodies against PLD1 and β-actin followed by infrared secondary antibodies. Detection was performed using LI-COR Odyssey imaging, and band intensities were normalized to β-actin using ImageJ. Detailed protocol in Supplemental Methods.

### Immunoprecipitation and Proteomic Analysis

PLD1-associated protein complexes were isolated from crude synaptoneurosomal fractions (BA38) of FTD and control brains using antibody-based immunoprecipitation (PLD1, CST #3832S) and magnetic bead capture. Complexes were validated by Western blot and subjected to LC–MS/MS analysis. Peptides were prepared using S-Trap digestion, analyzed on an Orbitrap Fusion mass spectrometer coupled to nanoLC, and searched against the human proteome using Proteome Discoverer with Sequest. Label-free quantification was performed without imputation. Detailed protocol in Supplemental Methods.

### Statistical Analysis

Data is represented as mean ± SEM. Group comparisons were performed using two-tailed Mann–Whitney U tests, with significance set at *P* < 0.05. Correlations were assessed using Pearson or Spearman coefficients as appropriate. All analyses were conducted in GraphPad Prism. Experiments and data analysis were double-blinded. Detailed protocol in Supplemental Methods.

## Results

### Functional synaptic integrity is reduced in BA38 and BA9 of FTD brains

Synaptic function was assessed using FASS-LTP in crude synaptoneurosomes from temporal (BA38) and frontal (BA9) cortices of FTD and control subjects (Table 1). Flow cytometry quantified GluA1–Nrx1β association as a measure of potentiation. In BA38, control samples exhibited significantly higher potentiation following chemical LTP compared to FTD (Fig. 1C–E; \*\*\**P* = 0.0003). A similar effect was observed in BA9 (Fig. 1F; \*\*\**P* = 0.0003). These findings indicate marked impairment of glutamatergic synaptic integrity in FTD across both regions.

**Fig-1:**
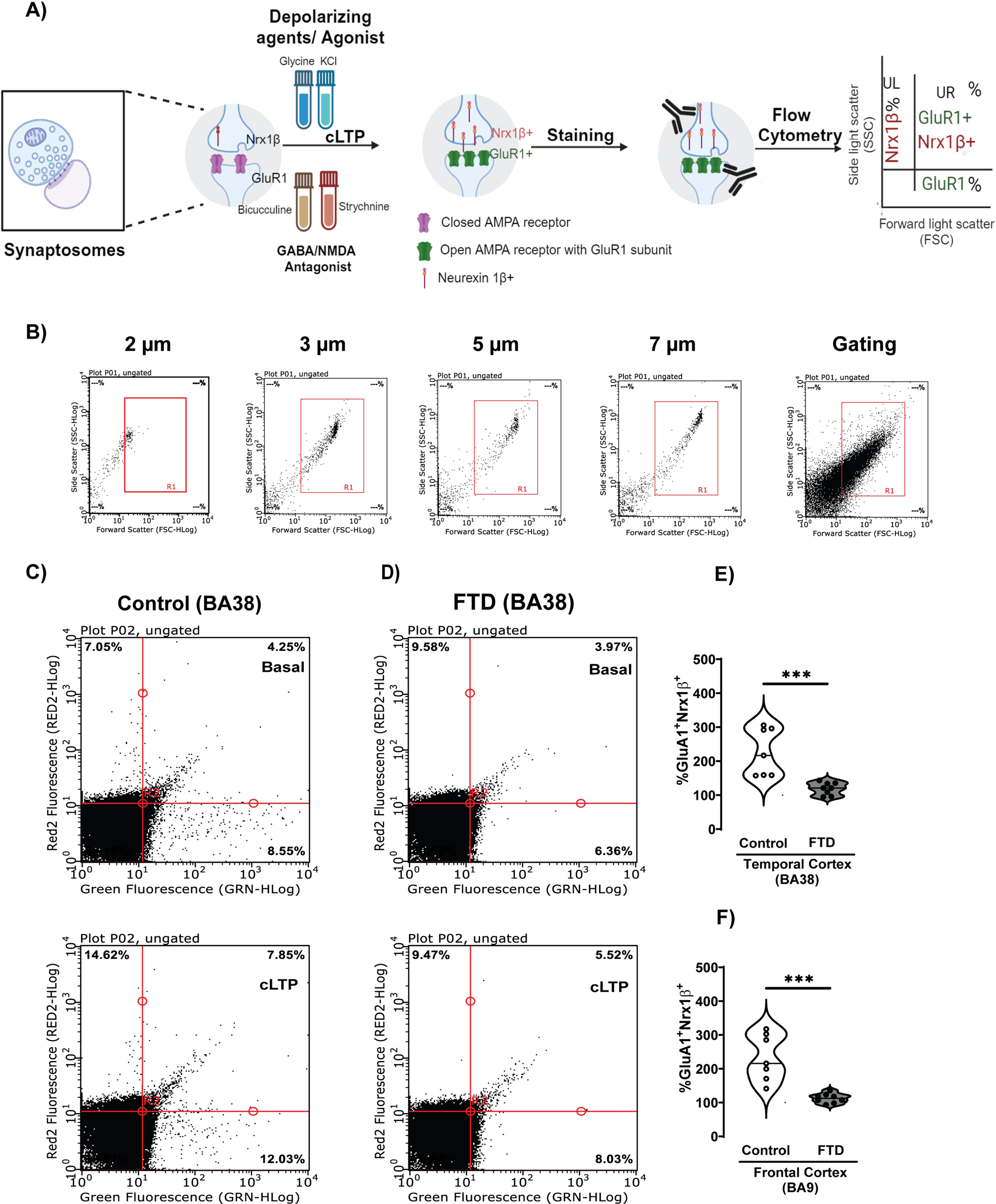
Glutamatergic neurotransmission is compromised in both BA38 and BA9 crude synaptoneurosomal fractions from FTD subjects compared to age-matched controls. (A) Steps involved in fluorescence-assisted synaptosome long-term potentiation (FASS-LTP) procedure. Crude synaptoneurosomes isolated from post-mortem human brains containing intact presynaptic and postsynaptic receptors are depolarized pharmacologically using agonists such as glycine and KCl, while simultaneously using antagonists such as strychnine and bicuculline for generating chemical LTP (cLTP). This elicits a response by increasing the number of surface GluA1 receptor subtypes available at the postsynaptic compartment, thereby facilitating the probability of presynaptic events. The percentage increase in the number of presynaptic and postsynaptic events is detected using respective antibodies, which are then captured in flow cytometry as double-positive events (GluA1^+^Nrx1β^+^) in the upper right (UR) quadrant. (B) Representative gating plots showing the size of the bead particles and crude synaptoneurosomes. Representative density plots from the temporal lobe (BA38) for (C) control and (D) FTD subjects showing GluA1-Nrx1β association being detected in the upper right quadrant following basal (top panels) and cLTP (bottom panels) treatment. Fluorescence thresholds were set by staining with secondary antibodies only (LL; lower left, UL; upper left quadrants). GluA1-Nrx1β increase in the UR was used as a measure of cLTP relative to basal levels. (E) Percent basal potentiation in BA38 for control subjects showed a higher percentage association of GluA1^+^Nrx1β^+^ compared to FTD patients; (control: 226.6 ± 26.46, FTD: 119.2 ± 6.417; \*\*\**P* = 0.0003). (F) Percent basal potentiation in BA9 for control subjects showed a higher percentage association of GluA1^+^Nrx1β^+^ compared to FTD patients; (control: 233.2 ± 26.04, FTD: 112.7 ± 4.895; \*\*\**P* = 0.0003). Values are expressed as mean ± SEM. *n*=7 (control), 8 (FTD), three technical repeats. Unpaired non-parametric Mann-Whitney test. Each circle in the graph represents a single subject.

### PLD1 expression is elevated in the temporal cortex but unchanged in the frontal cortex of FTD brains

Western blot analysis of crude synaptoneurosomal (P2) and cytosolic (S2) fractions revealed significantly higher PLD1 levels in BA38 of FTD compared to control subjects (crude synaptoneurosomal: \**P* = 0.0205; cytosolic: \*\**P* = 0.0022; Fig. 2A–B). In contrast, BA9 showed no significant differences in either fraction (Fig. 2C–D; *P* > 0.28). These findings indicate region-specific PLD1 upregulation in FTD and therefore we decided to investigate PLD1-tau interaction in the temporal (BA38) cortex in the current study.

**Fig-2:**
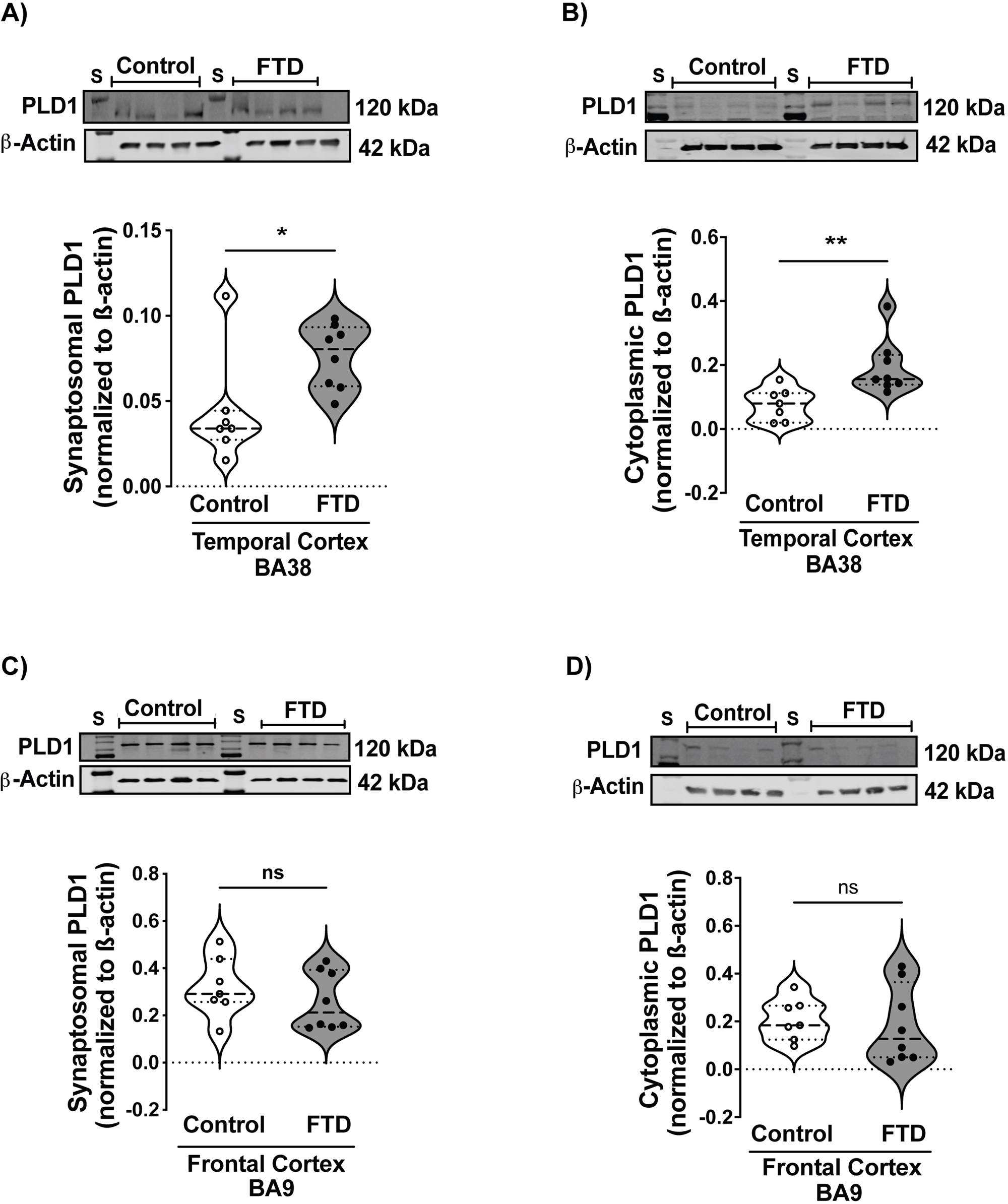
Crude synaptoneurosomal (P2) and cytoplasmic (S2) levels of PLD1 are differentially altered in frontal (BA9) and temporal (BA38) lobes of FTD patients compared to age-matched controls. Representative Western blot images of crude synaptoneurosomal and cytoplasmic fractions from control and FTD subjects. (A) Significantly elevated expression of PLD1 in crude synaptoneurosomal fractions of FTD patients compared to that of control subjects in BA38, \**P* = 0.0205, control: 0.04344 ± 0.01187, FTD: 0.07622 ± 0.006622. (B) Significantly elevated levels of PLD1 in cytoplasmic fractions of FTD patients compared to those of control subjects in BA38, \*\**P* = 0.0022, control: 0.07729 ± 0.01911, FTD: 0.1928 ± 0.03072. (C) No significant reduction in crude synaptoneurosomal expression levels of PLD1 in BA9 of FTD patients relative to the control subjects, *P* = 0.2810, control: 0.3431 ± 0.05052, FTD: 0.2609 ± 0.04358. (D) There is no significant change in PLD1 expression in cytoplasmic fractions of BA9 in FTD patients compared to that of the control subjects, *P* = 0.4634, control: 0.2070 ± 0.03273, FTD: 0.1844 ± 0.05681. S = Protein Standard. Values are expressed as mean ± SEM. *n*=7 (control), 8 (FTD). Three technical repeats were done for four subjects per group at a time.

### PLD1–tau interaction is strongly associated with pathological tau species in FTD temporal cortex

Bielschowsky staining revealed a significant increase in Pick bodies in BA38 of FTD brains compared to controls (\**P* = 0.0152), with a larger stained area (\*\**P* = 0.0022; Fig. 3A–B). Immunofluorescence showed robust PLD1 co-localization with total tau (HT7) in FTD (\*\**P* = 0.0012), alongside a trend toward elevated tau levels (*P* = 0.0721; Fig. 3D). PLD1 also significantly co-localized with hyperphosphorylated tau (AT8) (\*\**P* = 0.0022), which was increased in FTD (\**P* = 0.0401; Fig. 3E–F). Proximity ligation assay (PLA) confirmed PLD1–AT8 association (<40 nm) in FTD (\*\**P* = 0.0022; Fig. 3G–H). Furthermore, PLD1 exhibited strong co-localization with acetylated tau oligomers (TOMA2) in FTD (\*\*\**P* < 0.0001), which were markedly elevated compared to controls (\*\*\**P* < 0.0001; Fig. 3I–J). It is important to note that correlation analysis between phosphorylated (AT8) and non-phosphorylated tau (HT7) (Suppl. Fig. 1A) indicates no significant correlation, but when PLD1 colocalizes with each of them (Suppl. Fig. 1B) indicates significant correlation (r) at 0.7393, with a *P*-value is 0.0023, indicating significant correlation and a strong association of PLD1 with the pathological burden. These findings strongly implicate PLD1 in tau aggregation and neurotoxic tau species formation in FTD.

**Fig-3:**
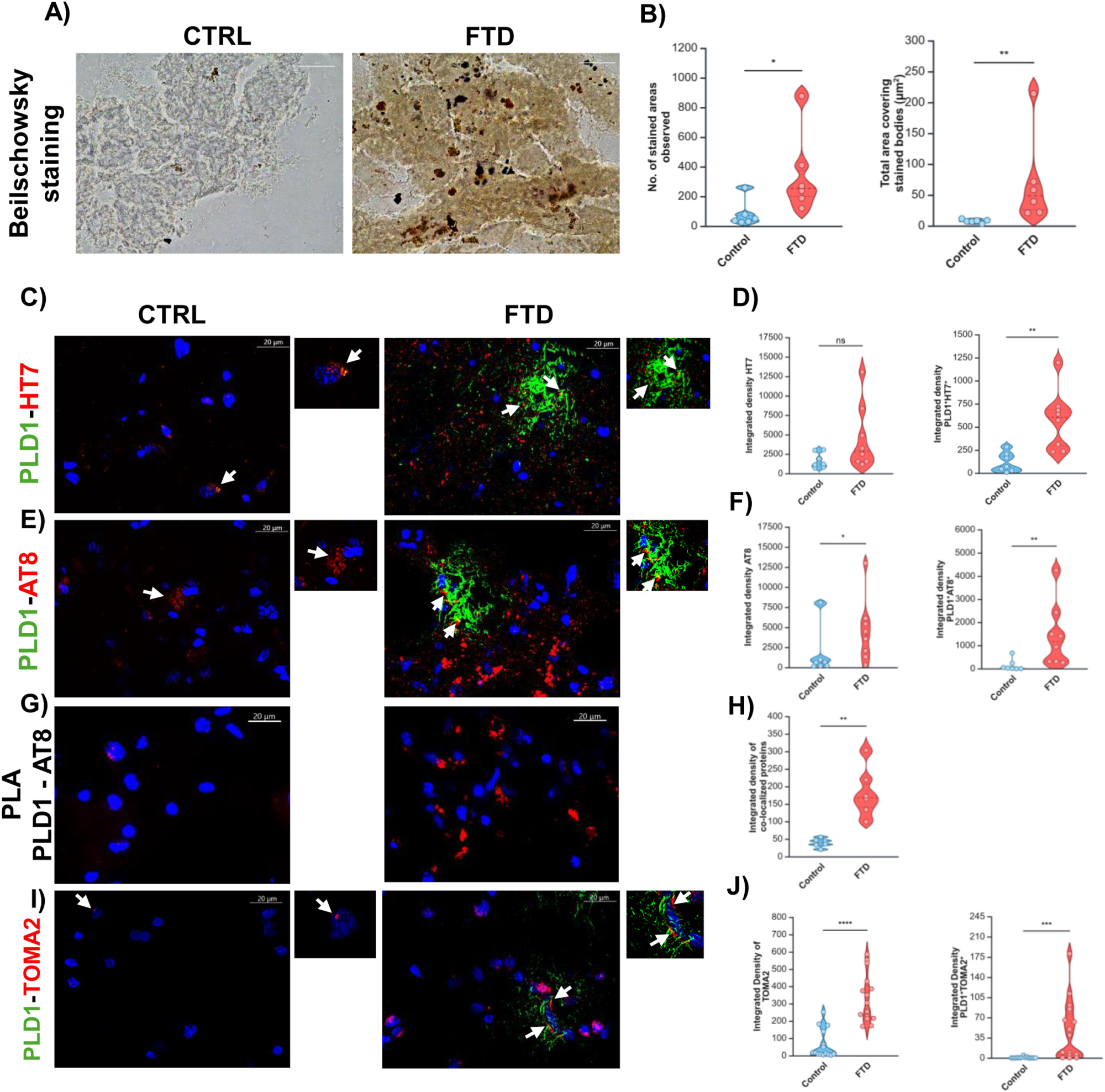
Enhanced Pick body formation and tau aggregation in the temporal lobe of FTD subjects linked to PLD1-tau interaction. (A) Representative bright-field microscopy images showing Bielschowsky silver-stained Pick bodies (black deposits) in the temporal lobe of control and FTD subjects. Scale bar = 20 µm. (B) Quantitative analysis of the number of stained areas (Pick bodies) revealed a significantly higher count in FTD subjects compared to controls (*\*P* = 0.0152; control: 80.18 ± 36.46, FTD: 350.3 ± 112.5). Quantitative analysis of the total area covered by the stained bodies showed a significantly increased area in FTD subjects compared to controls (*\*\*P* = 0.0022; control: 7.792 ± 1.136 µm², FTD: 71.01 ± 29.80 µm²). (C) Representative images showing double immunofluorescence of PLD1 (green) and total tau (HT7, red) in control and FTD subjects. (D) Quantitative analysis revealed a trend towards increased total tau (HT7) levels in FTD subjects, though not statistically significant (*P* = 0.0721; control: 1690 ± 372.7, FTD: 4551 ± 1477). However, a significant increase in PLD1 co-localization with HT7 was observed in FTD subjects (*\*\*P* = 0.0012; control: 114.9 ± 39.80, FTD: 573.5 ± 113.8). (E) Representative images showing double immunofluorescence of PLD1 (green) and hyperphosphorylated tau (AT8, red) in control and FTD subjects. (F) Quantitative analysis showed significantly elevated levels of hyperphosphorylated tau (AT8) in FTD subjects (*\*P* = 0.0401; control: 1565 ± 1094, FTD: 4538 ± 1402). A significant increase in PLD1 co-localization with AT8 was also observed in FTD subjects (*\*\*P* = 0.0022; control: 140.6 ± 93.67, FTD: 1425 ± 482.8). (G) Representative images showing PLA results for PLD1 and AT8 (red) in control and FTD subjects, indicating close proximity between the two proteins; DAPI (blue) stains nuclei. (H) Quantitative analysis of PLA images demonstrated a significant increase in PLD1-AT8 association within a 40 nm distance in FTD subjects (*\*\*P* = 0.0022; control: 39.11 ± 4.984, FTD: 182.1 ± 29.29). (I) Representative images showing double immunofluorescence of PLD1 (green) and acetylated tau oligomer (TOMA2, red) in control and FTD subjects. (J) Quantitative analysis revealed significantly elevated levels of tau oligomer (TOMA2) in FTD subjects (\*\**\*P* = <0.0001; control: 63.4413 ± 82.1798, FTD: 334.0428 ± 140.1164). A significant increase in PLD1 co-localization with TOMA2 was also observed in FTD subjects (*\*\*\*P* = <0.0001; control: 0.6427 ± 1.2517, FTD: 47.7100 ± 52.6851). Values are expressed as mean ± SEM. n = 6-7 (control), 6-8 (FTD). Unpaired non-parametric Mann-Whitney test. Scale bar = 20 µm. Each circle in the graph represents a single subject. Representative images were taken through the Keyence BZ-X810 microscope.

### PLD1 co-localization increases with astrocytes and shows distinct patterns with synaptic markers

Immunofluorescence revealed significantly elevated GFAP expression in BA38 of FTD compared to control brains (\**P* = 0.0270) and a robust increase in PLD1–GFAP co-localization (\*\*\**P* = 0.0003; Fig. 4A). These findings indicate astrocytic involvement in PLD1-associated pathology within the temporal cortex. Correlation analysis confirmed that these changes were not influenced by post-mortem interval (*P* > 0.13, Suppl. Fig. 1F). Differential patterns of PLD1 association with synaptic markers (Nrx1β, PSD95) were also observed, suggesting compartment-specific remodeling in FTD.

**Fig-4:**
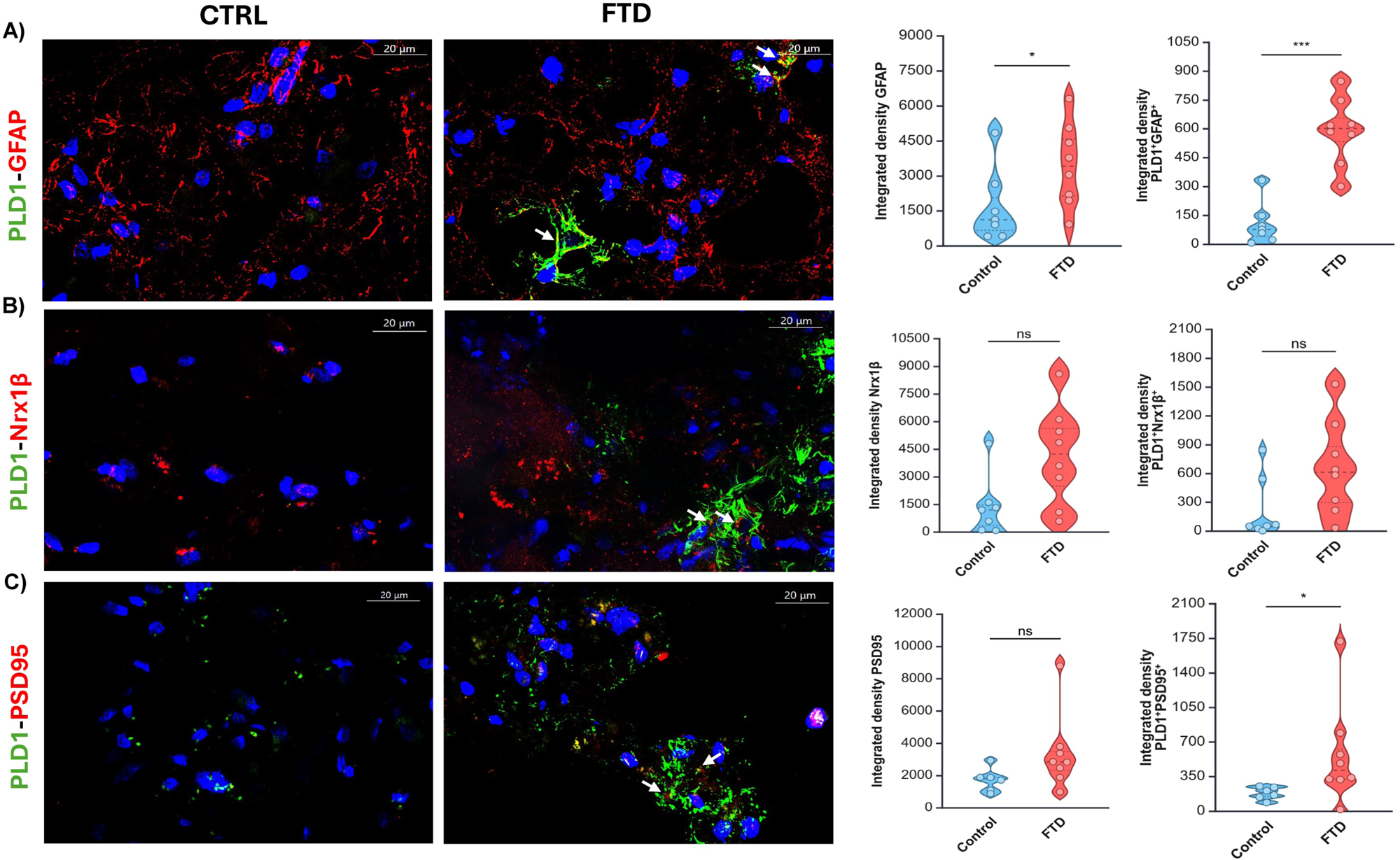
Region-Specific PLD1 Co-localization with Astrocytic and Synaptic Markers in the Brodmann Area 38. (**BA38) Region of FTD.** Representative immunofluorescence images and quantitative analyses from BA38 demonstrate enhanced co-localization of PLD1 with GFAP (astrocytic marker) in FTD compared to controls, indicating astrocytic involvement in PLD1-associated pathology. Additionally, differential increases in PLD1 co-localization with synaptic markers (e.g., Nrx1β, PSD-95) were observed in FTD brains, suggesting synaptic compartmentalization of PLD1 in disease states. (A) Representative double immunofluorescence images of PLD1 (green) and GFAP (red) in control and FTD subjects. Quantitative analysis revealed significantly elevated levels of GFAP in FTD compared to controls (*\*P* = 0.0270; control: 1695 ± 600.4, FTD: 3468 ± 629.9). Quantitative analysis demonstrated a significant increase in PLD1 co-localization with GFAP in FTD compared to controls (*\*\*\*P* = 0.0003; control: 106.9 ± 41.74, FTD: 589.6 ± 60.67). (B) Representative images showing double immunofluorescence of PLD1 (green) and Neurexin-1β (Nrx1β, red) in the temporal lobe of control and FTD subjects. Quantitative analysis revealed a trend but not significant increase in co-localization of PLD1 with Nrx1β in FTD (*P* = 0.0721; control: 227.9 ± 125.5, FTD: 656.7 ± 174). Nrx1β expression also showed a trend towards increase in FTD (*P* = 0.0541; control: 1395 ± 614.2, FTD: 4163 ± 942.2). (C) Representative images showing double immunofluorescence of PLD1 (green) and PSD95 (red) in control and FTD subjects. Quantitative analysis of PSD95 integrated density revealed no significant difference between control and FTD subjects (*P* = 0.2810; control: 2548 ± 501.5, FTD: 1186672 ± 1182150). Quantitative analysis demonstrated a significant increase in PLD1 co-localization with PSD95 in FTD compared to controls (*\*P* = 0.028; control: 372.8 ± 77.31, FTD: 237098 ± 235980). Values are expressed as mean ± SEM. Each circle represents an individual subject. Scale bar = 20 µm. Statistical significance was determined using an unpaired, non-parametric Mann-Whitney test.

### PLD1-associated proteomics reveals GFAP enrichment and synaptic pathway disruption in FTD

Targeted proteomics of PLD1 immunoprecipitates from BA38 synaptoneurosomes identified 28 significantly altered proteins (2-fold log change, *P* < 0.05; Fig. 5C–F). Comparative analysis revealed shift towards inflammatory and metabolic markers in the cytosol and depletion of vesicle-trafficking and proteostasis regulators in synaptoneurosomal fraction. GFAP showed a 4.45-fold increase in FTD (\**P* = 0.0361), corroborating immunofluorescence findings and reduction in presynaptic scaffolds and mitochondrial import components suggested impaired neurotransmission and energy homeostasisThese data support a PLD1-driven mechanism involving astroglial activation and synaptic proteostasis deficits in FTD.

**Fig-5:**
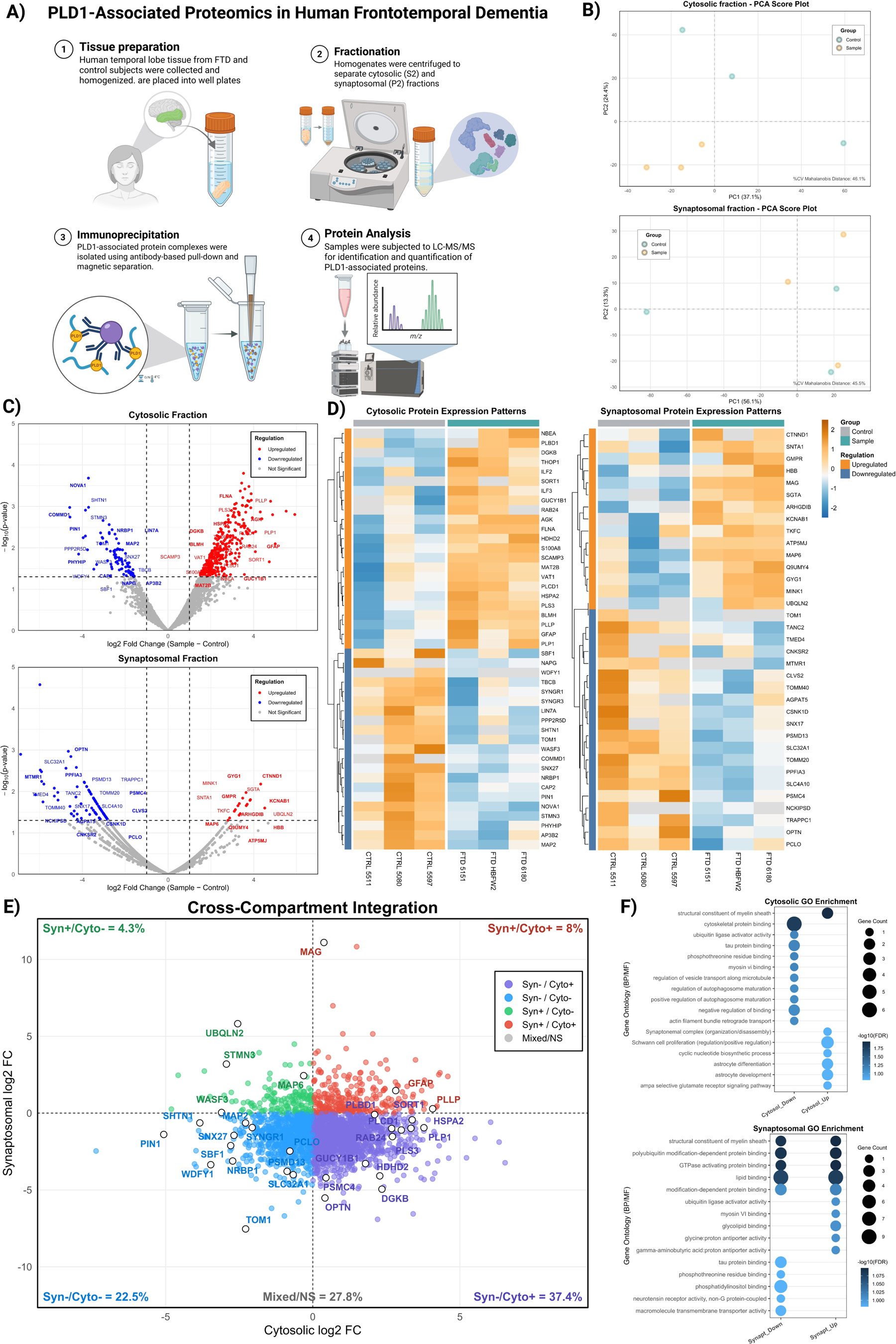
PLD1-associated proteomics in human frontotemporal dementia (FTD) reveals synaptic trafficking and proteostasis deficits accompanied by cytosolic metabolic, inflammatory, and astroglial remodeling. (A) Schematic of the experimental workflow. Human temporal lobe (BA38) tissues from FTD and control subjects were homogenized and fractionated to obtain cytosolic (S2) and crude synaptoneurosomal (P2) compartments. PLD1-associated protein complexes were isolated via antibody-based pull-down, validated by Western blotting, and subjected to LC–MS/MS for identification and quantification of PLD1-interacting proteins. (B) Principal component analysis (PCA) plots of cytosolic and crude synaptoneurosomal fractions show clear separation between FTD and control samples, indicating reproducible compartment-specific proteomic differences (Mahalanobis CV ≈ 45–46%). (C) Differential abundance analysis (volcano plots) highlights significant up- and down-regulated proteins in both compartments. In the cytosolic fraction, elevated levels of metabolic and lipid-signaling enzymes (DGKB, GUCY1B1, HDHD2), stress proteins (HSPA2), and GFAP, a canonical astrocytic marker, were observed, indicating increased glial and inflammatory activity. Decreased cytosolic PIN1 supports a permissive background for Tau misfolding. In the crude synaptoneurosomal fraction, presynaptic scaffolds (PCLO, PPFIA3), vesicle-trafficking regulators (SNX17, SLC32A1), and mitochondrial import components (TOMM20/40) were markedly reduced, suggesting synaptic vesicular and energy-transfer deficits. (D) Heatmaps of protein expression patterns illustrate coherent compartment-specific shifts. Cytosolic profiles exhibit upregulation of metabolic and glial-response modules, including GFAP, while crude synaptoneurosomal clusters show concerted downregulation of trafficking and proteostasis regulators (PSMC4, PSMD13, OPTN, SNX17). (E) Cross-compartment integration analysis maps cytosolic versus crude synaptoneurosomal log₂ fold changes, revealing that 37.4 % of proteins fall within the Syn^−^/Cyto^+^ quadrant—proteins reduced at synapses but increased in the cytosol—consistent with trafficking and degradative impairments. GFAP lies within this quadrant, reflecting redistribution or compensatory cytosolic upregulation associated with glial reactivity and neuronal stress. (F) Gene Ontology enrichment analysis shows that cytosolic changes are dominated by lipid metabolism, cyclic-nucleotide synthesis, cytoskeletal remodeling, and stress-response pathways (including astrocyte activation), whereas crude synaptoneurosomal alterations involve vesicle trafficking, proteasome regulation, and Tau- and phosphatidylinositol-binding proteins.

## Discussion

FTD, the second most common cause of dementia after AD, remains difficult to diagnose and dissect mechanistically[41]. Unlike AD, which primarily affects episodic memory, FTD manifests with early social disinhibition, motivational deficits, and behavioral changes, with cognitive decline emerging later[45]. The pronounced laminar and regional distribution of tau aggregates in FTD[42] disrupts long-range cortical connectivity, which underlies the associated behavioral and executive deficits[43,44]. Our study provides a comprehensive mechanistic framework linking synaptic dysfunction to tau pathology in FTD, with PLD1 as a central player in synaptic dysfunction.

Using our validated human FASS-LTP protocol, we first established robust functional synaptic deficits in glutamatergic neurotransmission within both BA38 and BA9, regions critically implicated in FTD (Fig 1). While prior studies have established glutamatergic dysregulation in FTD, these findings provide the first functional evidence for such deficits in human postmortem synaptic structures[46–48]. Importantly, PLD1 expression was significantly elevated in BA38 but not BA9 (Fig. 2), consistent with our prior observations of a brain region-specific pathology in a mouse model of AD/ADRD[19,49]. This differential pattern aligns with variability in BA9 synaptic profiles[50] and the consistent tau burden reported in BA38[2,7,51]. Although this observation provides the rationale for the concentration of our studies within BA38, further investigation will be required to elucidate the cell-type and subcellular compartment specific PLD1 splice variants[52–56] and isoforms[57–62] contributing to the PLD1-tau pathology in brain-region specific progression of FTD. In agreement, a recent report emphasizes similar complexities involving *C9orf72* splice variants in disease progression[63].

Following confirmation of tau pathology by Bielschowsky staining (Fig. 3), we established PLD1–tau interactions. Immunofluorescence revealed co-localization of PLD1 with both total and pathological tau species. AT8 detects phosphorylated tau at Ser202/Thr205 (and Ser208)[64], while TOMA2 is specific for astrocytic tau oligomers[65,66]. PLA provided first direct evidence of PLD1–tau molecular interaction, establishing a mechanistic link between PLD1 dysregulation and tau aggregation. These findings corroborate our prior work in AD/ADRD-like mouse models[20–22], where PLD1 attenuation restored dendritic spine integrity and rescued memory deficits in 3xTg-AD mice, implicating PLD1 in synaptic resilience. Our study in wildtype mice clearly demonstrated that PLD1 attenuation prevents TauO-driven synaptic dysfunction and memory deficit[19]. BA38 co-localization of PLD1 with multiple tau pathology markers thus suggests a mechanistic link between PLD1 activity and tau-driven neurodegeneration, likely accelerating synaptic dysfunction[12], and leading to progressive loss of excitatory neurons[43,67].

Beyond neuronal compartments, PLD1 was markedly enriched in astrocytes, as evidenced by PLD1–GFAP (Fig. 4) and TOMA2 co-localization (Fig. 3) along with its proteomic overrepresentation (Fig. 5). This astroglial involvement suggests that PLD1 contributes to neurodegeneration through both neuronal and non-neuronal pathways. This is consistent with emerging literature on glial mechanisms in FTD[4,5,68], where neuroinflammatory responses—such as astrocyte reactivity and T-cell infiltration—are associated with increased tau pathology that exacerbates neurodegeneration in BA38[69].

The PLD1-tau interaction likely drives the observed proteomic profile (Fig 5), characterized by the upregulation of metabolic, lipid-signaling, and stress-response proteins in the cytosol, concurrent with synaptic and mitochondrial deficits detected in the crude synaptoneurosomal fraction. This outcome is likely mediated by several converging mechanisms associated with tauopathy and neurodegeneration. Tau protein disruption in neurodegeneration modifies lipid composition and metabolism and promotes the upregulation of enzymes such as diacylglycerol kinase beta (DGKβ) and other lipid-signaling proteins[70,71]. These alterations are associated with subsequent increases in GFAP and heat shock proteins (e.g., HSPA2)[72,73]. PLD1 could exacerbate this collective set of processes[74]. Given that previous research, including our studies, indicate that PLD1 dysregulation impairs vesicle fusion and neurotransmitter release, resulting in synaptic loss[20,21,75–79], the synaptic and mitochondrial deficits identified here (Fig 5) are consistent with tau pathology-dependent progression in neurodegenerative disorders[80–84]. Specifically, these observed deficits include the disruption of presynaptic scaffold proteins (e.g., Piccolo, PPFIA3), vesicle-trafficking regulators (e.g., Sorting Nexin 17, SLC32A1), proteostasis factors (e.g., Pin1), and mitochondrial import components (e.g., TOMM20/40). Collectively, these findings support the hypothesis that the PLD1-tau interaction generates a compartmentalized effect. This is further evidenced by the upregulation of metabolic, lipid-signaling, and glial pathways in the cytosol, concurrent with the downregulation of trafficking and proteostasis regulators within the crude synaptoneurosomal fraction. Mechanistically, this differential impact suggests that the interaction drives metabolic and inflammatory stress in the cytosolic compartment, leading to synaptic and mitochondrial dysfunction at the synapse.

Future studies will focus on elucidating the specific mechanistic details of this complex PLD1-tau interactions. Such research is necessary to define the distinct contributions of both neuronal and non-neuronal pathways to the progression of FTD.

## Supporting information

Supplemental Table and Detailed Methods

## Declarations

### Ethics Approval and Consent to Participate

Postmortem, de-identified brain tissues used in this study were obtained from the NeuroBioBank (NBB) Maryland, Baltimore, through a material transfer agreement. Brain donors were enrolled with NBB and underwent neuropathological evaluation postmortem, in accordance with the institutional review board of the University of Miami Brain Endowment Bank.

### Consent for Publication

Not applicable.

### Availability of Data and Materials

All data generated or analyzed during this study are included in this published article and its supplementary information files.

## Acknowledgements

Brain tissue samples were obtained from the National Institutes of Health (NIH) NeuroBioBank with the MTA number NBB1001 and 1019. The authors express their gratitude to the brain donors and their families. The authors thank Anusha Srinivasan for her critical review of the manuscript. The authors are thankful to Dr. Rakez Kayed for providing us with their in-house TOMA2 antibody for the studies and would also like to thank Dr. William Russell and the Mass Spectrometry Core facility for helping us with the procedural specifics for proteomics study. The authors are grateful to Dr. Giulio Taglialatela and his lab for their constructive feedback and acknowledge the Mitchell Center for Neurodegenerative Diseases for providing shared resources.

## Authors’ Contributions

S.B., C.N., S.G.S.S., and B.K. conceived and designed the study, and wrote the manuscript. S.B., C.N., S.G.S.S., P.C., B.T., and B.K. performed the experiments and analyzed the data. S.B., S.G.S.S., C.N., and B.K. analyzed and interpreted the data. C.N. conducted proteomics, Western blots, and some immunofluorescence experiments. S.B. performed immunofluorescence, proximity ligation assays, and Bielschowsky staining. B.T. and C.N. conducted FASS-LTP experiments. S.B. performed Pearson co-localization analysis of immunofluorescence data. L.K. contributed to interpretations for proteomics. M.M. and G.T. provided intellectual input on proximity ligation assay-based approaches. All authors read and approved of the final manuscript.

