## Supplemental Table and Detailed Methods for "Exploring the PLD1-tau interaction in Frontotemporal Dementia"

**Supplementary Figures**

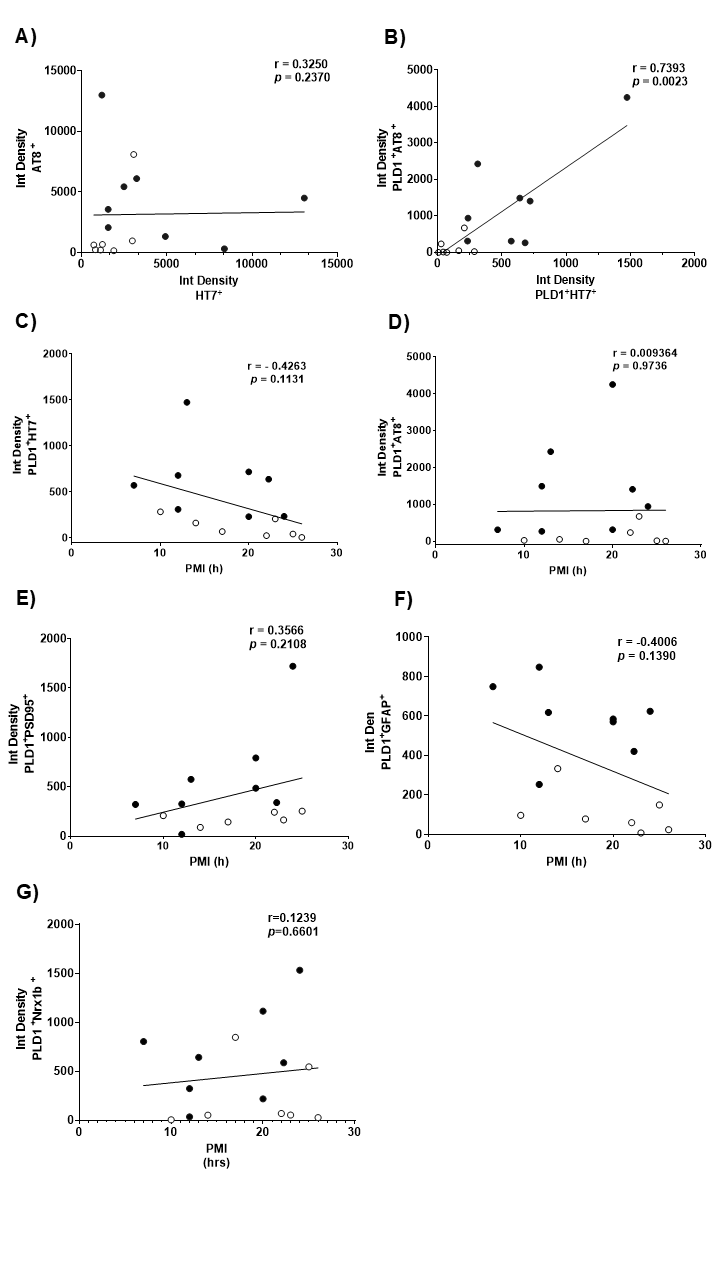
s

**Suppl. Fig-1: Post-Mortem Interval (PMI) Correlation Analysis in the BA38 Region of Control and FTD Brains.** This figure presents Pearson correlation analyses to assess the potential influence of post-mortem interval (PMI) on the integrated density measurements of various protein interactions and individual protein levels in the BA38 region (temporal lobe) of control and frontotemporal dementia (FTD) brain tissues. (A) Correlation analysis between phosphorylated (AT8) and non-phosphorylated tau (HT7). The correlation coefficient (r) is 0.3250, and the *P*-value is 0.2370, indicating no significant correlation (ns). (B) Correlation analysis between phosphorylated (AT8) and non-phosphorylated tau (HT7) when PLD1 colocalizes with each of them. The correlation coefficient (r) is 0.7393, and the *P*-value is 0.0023, indicating significant correlation and a strong association of PLD1 with the pathological burden. (C) Correlation analysis between the integrated density of PLD1 co-localized with total tau (HT7) and PMI. The correlation coefficient (r) is -0.4263, and the *P*-value is 0.1131, indicating no significant correlation (ns). (D) Correlation analysis between the integrated density of PLD1 co-localized with phosphorylated tau (AT8) and PMI. The correlation coefficient (r) is 0.009364, and the *P*-value is 0.9736, indicating no significant correlation (ns). (E) Correlation analysis between the integrated density of PLD1 co-localized with PSD95 and PMI. The correlation coefficient (r) is 0.3566, and the *P*-value is 0.2108, indicating no significant correlation (ns). (F) Correlation analysis between the integrated density of PLD1 co-localized with GFAP and PMI. The correlation coefficient (r) is -0.4006, and the *P*-value is 0.1390, indicating no significant correlation (ns). (G) Correlation analysis between the integrated density of PLD1 co-localized with Nrx1β and PMI. The correlation coefficient (r) is 0.1239, and the *P*-value is 0.6601, indicating no significant correlation (ns). Each data point in the graphs represents an individual subject. White-filled circles represent control subjects, and black-filled circles represent FTD subjects. See Table 1 for individual subject IDs and their corresponding PMI values. Statistical significance was determined using an unpaired, non-parametric Mann-Whitney test.

**Supplementary Methods:**

**Human Subjects and Tissue Processing**

Postmortem brain tissues were obtained from the NIH NeuroBioBank (Maryland, USA). Fresh-frozen temporal cortex blocks (BA38; *n* = 8 per group) were stored at –80 °C, equilibrated at –20 °C, and embedded in optimal cutting temperature (O.C.T.) compound (Tissue-Tek, #4582). Sections (12–16 µm) were cut using a cryostat and mounted on Superfrost Plus slides (Thermo Fisher Scientific, #12-550-15). Slides were stored at –20 °C until use.

**Fixation and Blocking:** Sections were fixed in 4% paraformaldehyde (0.1 M PBS, pH 7.4) for 30 min at room temperature (RT). Non-specific binding was blocked with 5% bovine serum albumin (BSA; Thermo Fisher Scientific, #A4503-50G) and 10% normal goat serum (NGS; Millipore Sigma, #S26-50 mL). Permeabilization was performed using 0.5% Triton X-100 and 0.05% Tween-20 for 1 h at RT.

**Primary Antibody Incubation:** Slides were incubated overnight at 4 °C with primary antibodies diluted in PBS containing 1.5% NGS. Antibodies included:

- Rabbit anti-PLD1 (1:200; Abcam, #ab50695; RRID: AB_2237051)
- Mouse anti-AT8 (1:200; Invitrogen, #MN1020)
- Mouse anti-tau HT7 (1:2000; Thermo Fisher Scientific, #MN1000; RRID: AB_2314654)
- Mouse anti-neurexin (1:200; NeuroMab, #N170A)
- Mouse anti-PSD95 (1:200; Abcam, #13552; RRID: AB_300453)
- Chicken anti-GFAP (1:200; Aves, #GFAP5727980)

**Secondary Antibody Incubation:** After three PBS washes (10 min each), slides were incubated for 1 h at RT with Alexa Fluor-conjugated secondary antibodies diluted in PBS with 1.5% NGS:

- Goat anti-rabbit Alexa Fluor 488 (1:400; Invitrogen, #A11034; RRID: AB_2576217)
- Goat anti-mouse Alexa Fluor 594 (1:400; Invitrogen, #A11032; RRID: AB_2534091)
- Goat anti-chicken Alexa Fluor 594 (1:400; Invitrogen, #A11039; RRID: AB_2534099)

**Autofluorescence Quenching and Mounting:** Slides were treated with 0.3% Sudan Black in 70% ethanol for 10 min to reduce lipofuscin autofluorescence, rinsed in deionized water, and mounted using Fluoromount-G containing DAPI (SouthernBiotech, #0100-20).

**Exclusions:** Control subject 5890 (PMI = 37 h) was excluded from all analyses; subject 5862 was excluded from PLD1–PSD95 co-localization due to limited tissue availability.

Images were acquired using a Keyence BZ-X810 microscope under identical exposure settings for all groups.

**FASS-LTP Protocol**

Frontal and temporal postmortem brain tissues were homogenized using the Syn-PER extraction method [8,16,17]. FASS-LTP was conducted according to established protocols [16,18,19].

**Buffer Preparation:**

- **Extracellular solution (mM):** 120 NaCl, 3 KCl, 2 CaCl₂, 2 MgCl₂, 15 glucose, 15 HEPES, pH 7.4
- **cLTP solution (mM):** 150 NaCl, 2 CaCl₂, 5 KCl, 10 HEPES, 30 glucose, pH 7.4 (Mg²⁺ omitted)

**Procedure:**

1. Synaptosomal P2 fractions were suspended in 200 µL of extracellular (E), basal (B), or cLTP (C) solution and incubated at RT for 30 min to recover and activate synaptoneurosomes.
2. Bicuculline methiodide (0.02 mM) and strychnine (0.001 mM) were added to tube C only. Glycine (5 mM; 20 µL) was added to tube C as an NMDAR co-agonist; extracellular solution was added to tubes E and B.
3. Depolarization was induced by adding 100 µL of KCl solution (50 mM NaCl, 100 mM KCl, 2 mM CaCl₂, 30 mM glucose, 10 mM HEPES, 0.5 mM glycine, 0.001 mM strychnine, 0.02 mM bicuculline) to tube C and incubating at 37 °C for 30 min.
4. Reactions were stopped by adding 0.5 mL of 0.1 mM EDTA-PBS and 4 mL of 5% blocking buffer, followed by centrifugation (2500 g, 5 min, 4 °C).
5. Pellets were incubated with primary antibodies: GluA1 (rabbit polyclonal, ABN241, EMD Millipore) and Nrx1β (mouse monoclonal, N170A/1, Antibodies Inc) at 2.5 µg/mL for B and C tubes; E tube received blocking buffer only.
6. After two washes, pellets were resuspended in secondary antibody solution (anti-rabbit Alexa Fluor 488 and anti-mouse Alexa Fluor 647, Invitrogen) and incubated for 45 min at 37 °C, protected from light.
7. Samples were fixed in 2% paraformaldehyde and analyzed using a Guava easyCyte™ 8 flow cytometer (GuavaSoft 2.7).

Double-positive events (GluA1⁺Nrx1β⁺) in the upper-right quadrant were quantified as a measure of cLTP relative to basal levels.

**Supplementary Methods: Bielschowsky Staining Protocol**

Bielschowsky silver staining was performed to detect tau deposits and Pick bodies, characteristic of FTD, following an established protocol [20] with minor modifications.

**Procedure:**

1. Cryosections (12–16 µm) were stored at –20 °C until use.
2. Slides were hydrated in distilled water for 5 min, repeated three times.
3. A Coplin jar containing 20% silver nitrate was preheated in a microwave for 35 s; slides were incubated at 37 °C for 15 min and washed in distilled water (3 × 5 min).
4. Ammonium hydroxide was added dropwise to the silver nitrate solution with stirring until precipitates dissolved; slides were incubated at 37 °C for 10 min.
5. Developer solution (formaldehyde, citric acid, nitric acid) was prepared fresh; slides were immersed for 1–5 min until tissue stained black.
6. Slides were rinsed in ammonia water for 2 min, followed by three washes in distilled water (2 min each).
7. Excess stain was removed by immersing slides in 5% Hypo solution for 2 min, followed by three washes in distilled water (3 min each).
8. Slides were dehydrated through graded ethanol, cleared, and mounted with coverslips.

**Note:** All solutions were freshly prepared. Six subjects per group (control and FTD) were processed under identical conditions.

**Quantitative Analysis**

Brightfield microscopy was performed using a Keyence BZ-X810 microscope. For each subject, five randomly selected regions of interest were imaged. Quantitative analysis was conducted using Keyence Analyzer software. Briefly, images were imported into the software, and areas containing Bielschowsky silver-stained Pick bodies were manually delineated using the polygonal selection tool. Transparency settings were adjusted to optimize visualization of stained deposits. The software then computed two metrics: (i) the number of discrete Pick body-positive regions and (ii) the total area occupied by these regions (expressed in μm²). These values were aggregated across five fields per subject for statistical analysis.

**Proximity Ligation Assay (PLA)**

Six control and six FTD tissue samples were fixed in 4% paraformaldehyde for 30 min at RT, washed thrice in distilled water (10 min each), and blocked in Duolink® blocking solution at 37 °C for 1 h. Sections were incubated overnight at 4 °C with rabbit anti-PLD1 (1:200; Abcam, ab50695, RRID: AB_2237051) and mouse anti-AT8 (1:100; Invitrogen, MN1020, RRID: AB_223647) diluted in Duolink® antibody diluent. After two washes in 1× Wash Buffer A, PLUS and MINUS PLA probes were applied (1:5 ratio) for 1 h at 37 °C. Ligation was performed using ligation buffer (1:5) and ligase (1:40) for 30 min at 37 °C, followed by amplification with amplification buffer (1:5) and polymerase (1:80) for 100 min at 37 °C. Slides were washed twice in 1× Wash Buffer B (10 min each) and once in 0.01× Wash Buffer B (1 min), then mounted with Duolink® in situ mounting medium containing DAPI. Images were acquired using a Keyence BZ-X810 microscope.

**Quantitative microscopy**

Images for immunofluorescence and PLA were acquired using a Keyence BZ-X810 microscope with 10X, 60X, and/or 100X objectives. For each subject, two sections were analyzed, and five images per section were captured. Quantitative analysis was performed using ImageJ software (NIH), and the fluorescence intensity of each marker was analyzed as integrated density (IntDens). For PLA, a red, bright signal, indicating two proteins within a <40 nm distance, was generated and quantified as fluorescence intensity.

**Tissue Processing and Western Blot Analyses**

Frozen tissue blocks were processed in Syn-PER buffer (catalog #87793) with 1% protease/phosphatase inhibitor cocktail (catalog #P8430, Sigma-Aldrich). Homogenates were centrifuged at 1,200 × g for 10 min at 4°C; supernatants were centrifuged at 15,000 × g for 20 min at 4°C. Pellets were resuspended in HEPES-buffered Krebs-like solution (143.3 mM NaCl, 1.2 mM CaCl₂, 4.75 mM KCl, 1.3 mM MgSO₄·7H₂O, 0.1 mM NaH₂PO₄, 20.1 mM HEPES, 10.3 mM D-glucose; pH 7.4). Protein concentrations were measured using Nanodrop 2000C (Thermo Scientific).

Proteins were separated by SDS-PAGE and transferred to nitrocellulose membranes (catalog #10600001, Sigma-Aldrich) at 100 V for 35 min at 4°C. Membranes were blocked in Odyssey buffer (LI-COR) for 1 h at RT and incubated overnight at 4°C with primary antibodies PLD1 (1:500, CST), β-actin (1:50,000, Sigma-Aldrich). After TBST washes, membranes were incubated with IRDye® secondary antibodies (LI-COR) at 1:10,000 dilution for 1 h. Imaging was performed using Odyssey Infrared Imaging System (LI-COR). Band densities were analyzed using Image Studio Lite and ImageJ, normalized to β-actin. Representative images were compiled in TIFF format using Microsoft PowerPoint.

**Immunoprecipitation and Proteomics Workflow**

**Immunoprecipitation:** P2 fractions (40 µg) were incubated overnight with PLD1 antibody (0.2 µg) at 4°C, followed by conjugation with Dynabeads M-270 epoxy slurry (60 µl; 3 h). Pellets were washed thrice in Tris buffer (pH 7.5), verified by Western blot, and processed for LC–MS/MS.

**Sample Digestion:** Proteins were solubilized in SDS/TEAB, reduced (TCEP), alkylated (iodoacetamide), and quenched (DTT). After acidification and methanol-based binding, samples were loaded onto S-Trap spin columns, washed, and digested with trypsin (1:25 ratio, 4 h at 37°C). Peptides were eluted sequentially with TEAB, formic acid, and acetonitrile, dried, and resuspended for LC-MS/MS.

**NanoLC-MS/MS:** Peptides were analyzed using an UltiMate 3000 RSLCnano system coupled to an Orbitrap Fusion mass spectrometer. Gradient elution (2–90% ACN over 120 min) was performed on C18 columns. MS acquisition used top-speed DDA with Orbitrap survey scans (120,000 resolution) and CID fragmentation in the ion trap. Dynamic exclusion was applied (±10 ppm, 15 s).

**Proteomics Data Analysis:** All downstream proteomic analyses were performed in R, with differential expression between FTD and control samples quantified using limma eBayes and initially assessed using FDR-adjusted p-values. As the regular stringent correction parameters identified very few detectable proteins in this dataset, the plots were generated using supplementary thresholds (|log2FC| ≥ 1 with raw p < 0.05 for volcano and heatmap analyses, |log2FC| ≥ 0.4 with FDR < 0.20 for compartmental integration) to illustrate broader effect-size patterns within the data. Gene labels in volcano plots were restricted to biologically relevant markers to maintain clarity and interpretability. PCA was performed using prcomp on centered and scaled intensities, with variance explained by PC1 and PC2 reported for each fraction, and Mahalanobis distance based %CV used to evaluate sample-level variance structure. Functional enrichment analyses were conducted using gProfiler2 with Benjamini–Hochberg FDR correction (FDR < 0.20) and visualized as GO bubble plots.

**Statistical Analysis**

All statistical analyses were performed using GraphPad Prism (v9 or v10; San Diego, CA, USA). Non-parametric tests (Mann–Whitney U or Wilcoxon rank-sum) were applied to account for non-normal data distributions. Correlations between experimental parameters and PMI were evaluated using Pearson correlation coefficients, while Spearman correlation was used for co-localization metrics. Significance threshold was set at *P* < 0.05. Double blinding was implemented by coding samples prior to experimentation and analysis; codes were broken only after completion of data analysis.
